# Inbreeding depression and population viability in a recovering population of Mauritius kestrels

**DOI:** 10.64898/2026.07.30.741698

**Authors:** Ken Norris, Carl G. Jones, Jim Groombridge, Sion Henshaw, Hernán E. Morales, Kevin Ruhomaun, Vikash Tatayah, Cock van Oosterhout, Xuejing Wang, Nicolas Zuel, Malcolm A.C. Nicoll

## Abstract

Inbreeding depression (the reduction in fitness associated with inbreeding) has been demonstrated in a wide range of animals, but despite its ubiquity, is not an inevitable consequence of inbreeding. As a result, there is uncertainty about the extent to which inbreeding depression poses an ongoing risk to endangered species currently experiencing significant demographic recovery. Quantifying inbreeding depression will be critical if we want to understand these risks. A comprehensive quantification of the fitness costs of inbreeding requires detailed individual-based longitudinal data so lifetime impacts can be assessed. Here, we use an extraordinarily detailed long-term dataset on Mauritius kestrels (*Falco punctatus*) to explore inbreeding depression in a population currently experiencing significant demographic recovery. To do so, we constructed a social pedigree of 1,758 individuals and combined this with 1,240 nest records and 1,411 individual resighting histories to explore lifetime fitness effects over a 30-year period. Inbreeding increased significantly over time as the population recovered before stabilising. Inbred eggs were less likely to survive to fledging. Inbred adult male and female birds had significantly lower annual reproductive success than outbred individuals because of a lower annual egg-to-fledgling survival probability. This resulted in significantly lower lifetime reproductive success in inbred females but not males, which showed a negative trend. Population growth was negative and extinction risk increased slightly at current levels of inbreeding. These impacts will become more severe should inbreeding levels increase in the future, which is highly likely given ongoing genomic erosion. Taken together, our results demonstrate significant fitness costs associated with inbreeding in Mauritius kestrels, which pose an ongoing risk to population viability. This suggests that monitoring and managing inbreeding risks in endangered species will likely be required even in populations that are showing significant demographic recovery in response to conservation interventions.

## Introduction

Inbreeding depression (the reduction in fitness associated with inbreeding) is common in birds and mammals (Keller, 1998, Crnokrak and Roff, 1999, Slate et al., 2000, Keller and Waller, 2002, Szulkin et al., 2007, Wells et al., 2018, Leedale et al., 2020, Hasselgren et al., 2021, Sin et al., 2021, Beccardi et al., 2024), and widely regarded as a significant risk to the viability of endangered species (Armstrong et al., 2021, Kennedy et al., 2014, Harrisson et al., 2019, Trask et al., 2021, Hasselgren et al., 2021). Despite its ubiquity, inbreeding depression is not an inevitable consequence of inbreeding. This is because it depends on the extent to which purging selection removes deleterious, recessive variants unmasked by small population size (Hedrick and Garcia-Dorado, 2016, Dussex et al., 2021, Dussex et al., 2023). If purging removes most of these variants, then inbreeding may still occur but with limited fitness consequences (Robinson et al., 2018, Robinson et al., 2022). In contrast, if purging is incomplete, populations may still be at risk from inbreeding depression, which could reduce population viability in the longer-term (Kennedy et al., 2014, Bozzuto et al., 2019, Harrisson et al., 2019, Robinson et al., 2019, Grossen et al., 2020, Armstrong et al., 2021, Trask et al., 2021). This has important implications for endangered species because it is currently unclear whether inbreeding depression poses an ongoing risk to populations currently experiencing significant demographic recovery (Femerling et al., 2023, Fontsere et al., 2025). Distinguishing between these alternatives requires us to better understand the fitness costs of inbreeding in recovering populations and their consequences for population viability.

Quantifying the fitness costs of inbreeding is data demanding. A comprehensive assessment of inbreeding depression requires studies involving the lifetimes of multiple individuals experiencing a range of environmental conditions (Keller, 1998, Keller and Waller, 2002, Szulkin et al., 2007, Trask et al., 2021). This is because fitness costs might only be apparent during particular life stages, or be amplified or obscured by environmental conditions (Keller and Waller, 2002, Richardson et al., 2004, Szulkin and Sheldon, 2007, Kubacka et al., 2025). Furthermore, it is important to understand the implications of inbreeding depression for population viability in the longer-term (Armstrong et al., 2021, Trask et al., 2021). This type of integrated approach is needed to comprehensively assess the risks inbreeding depression poses to population recovery in endangered species.

Here, we take advantage of an extraordinarily detailed longitudinal study of an endangered wild bird, the Mauritius kestrel (*Falco punctatus*). Following a severe bottleneck in the 1980s (Groombridge et al., 2001), a population was re-established in the Bambou Mountains in the east of Mauritius through captive rearing and release (Jones et al., 1995, Nicoll et al., 2021). This population has subsequently grown and stabilised at ∼50 breeding pairs (Nicoll et al., 2021). During this period, the intensive monitoring of nesting attempts and individual mark-resighting studies have generated a remarkably complete dataset across the lifetimes of multiple individuals (details in the **Methods** section). This represents an ideal case study for asking whether inbreeding depression remains important in a population experiencing significant demographic recovery.

Our aim in this paper is to use these data to quantify the lifetime impact of inbreeding on fitness components in Mauritius kestrels and explore its consequences for population growth and viability (extinction risk). Previous studies have documented the loss of genetic variation through the population bottleneck (Groombridge et al., 2000), and recorded very high levels of inbreeding in the recovering population (Ewing et al., 2008). Furthermore, recent genomic analyses suggest significant and ongoing genomic erosion with incomplete purging of potentially deleterious variants (Wang et al., 2026).

Taken together, these findings increase the likelihood that deleterious, recessive mutations might be unmasked and cause significant inbreeding depression in this species. As a result, this is an issue of ongoing conservation concern.

## Methods

### Long-term conservation and monitoring programme

The Mauritius kestrel is a small falcon endemic to the forests of Mauritius, an island in the SW Indian Ocean. It is an iconic conservation success story. The wild population was reduced to only four known individuals in 1974 occurring in a single relict population in the Black River Gorges before intensive conservation management, including captive rearing and release, nest box provision, supplementary feeding and predator control helped the population recover. By 2018, the species occurred in three isolated populations and numbered roughly 140-170 mature individuals, although the Black River Gorges population has declined in recent years (Nicoll et al., 2021).

The population in the Bambou Mountains in the East of Mauritius was extirpated in the 1950s. Captive rearing and release in the late 1980s/early 1990s began to re-establish the population, and combined with the provision of nest boxes and predator control, the population grew then stabilised at ∼50 breeding pairs (Nicoll et al., 2021). Since its inception, this population has been intensively monitored. Released individuals were marked with a numbered metal ring and a unique combination of coloured leg rings.

Once released individuals began breeding, a basic monitoring programme was established that has been followed continuously since the early 1990s (Nicoll et al., 2021). During each breeding season, breeding pairs are located and the adult birds identified. Their nesting attempts are followed to establish first egg date, clutch size and the number of chicks fledged. Chicks are ringed in the nest with a numbered metal ring and a unique combination of coloured leg rings. This enables them to be subsequently identified if they are recruited into the breeding population. In this way, a detailed picture of the life histories of individual Mauritius kestrels has been developed. This long-term monitoring programme has generated two primary datasets – records of each nesting attempt (first egg date, clutch size and the number of chicks fledged, including the IDs of parents and offspring (nest records data); and a dataset containing mark-resighting histories for all marked individuals (resighting data).

### Data analyses

These longitudinal data are ideal for a comprehensive study of inbreeding depression across the lifetimes of individual birds (Fig. 1). Our approach treats an individual’s level of inbreeding (as measured by the inbreeding (*F*) coefficient) as a trait, and relates this individual trait to a series of fitness-related demographic traits that represent the major life stage transitions in a Mauritius kestrel life cycle (see Fig. 1). This includes lifetime reproductive success (LRS), which provides a short-term proxy for fitness.

**Figure 1.**
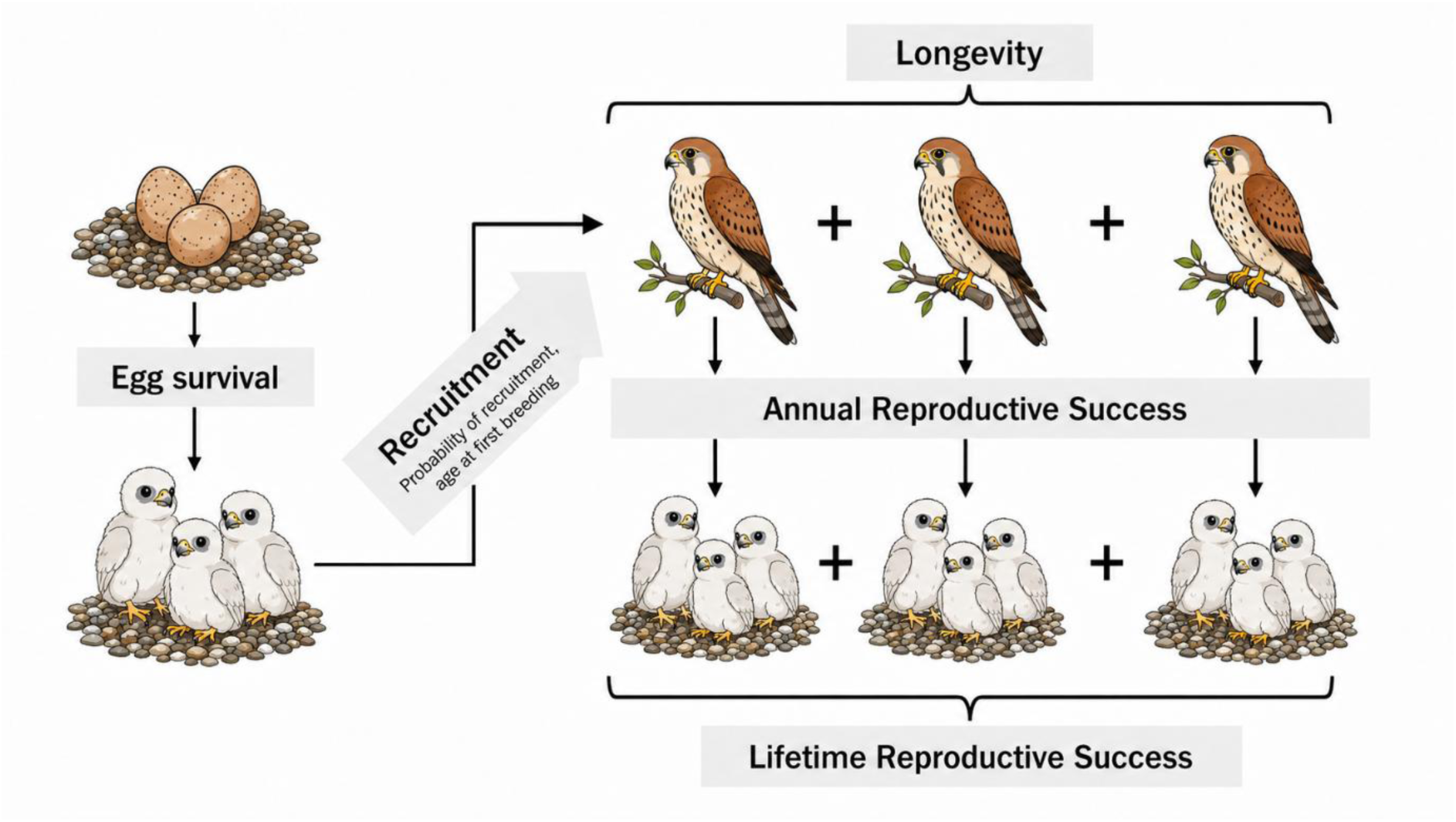
Diagram showing the life cycle of a Mauritius kestrel. Key stages are shown together with the demographic traits associated with those stages that formed the basis of our analyses.

For our analyses, we first used the parent-offspring data from the nest records to construct a social pedigree following the standard format – individual ID, mother (dame) ID, father (sire) ID. On a few occasions, an individual was first marked as a breeding adult. In these cases, we created a birth event for each individual assuming it was hatched in the previous breeding season to unknown parents. In this way, we ensured the pedigree conformed to the key assumption that an individual must first be born into the pedigree before appearing as a parent. Missing IDs were included in the pedigree as NAs. The resultant social pedigree consisted of 1,758 individuals.

The quality of a social pedigree is dependent on the completeness of individual identifications, so we calculated the percentage of adult female, adult male and breeding pairs identified over time from nest record data (Supplementary information; Fig. S1). This showed the percentage declined gradually as the population grew, but declined significantly through the Covid-19 pandemic before beginning to recover in recent years. As a result, we limited our analyses to the 1987/88-2017/18 breeding seasons to ensure the pedigree we used in our analyses was as complete as possible with the exception of the 2018/19 resightings data, which we used to censor individuals that remained alive beyond the main study window. Note that Mauritius kestrel breeding seasons span the austral spring and summer and so include two calendar years (e.g. 2009/10). We used the R package pedigree to calculate inbreeding (*F*) coefficients and pedigree depth for all individuals in this pedigree (Coster, 2022), and selected only those individuals with a pedigree depth ≥2 generations for further analyses (Keller, 1998). This resulted in 1,055 individuals with pedigree data available for downstream analyses.

For our analyses, we included nest records up to and including the 2017/18 breeding season to align with our pedigree. Following previous studies, we excluded any nests that had experienced conservation management (e.g. eggs removed, fostered chicks added) (Burgess et al., 2011, Burgess et al., 2008, Cartwright et al., 2014b, Cartwright et al., 2014a, Nevoux et al., 2013). This resulted in 1,240 nests available for analysis. From the resightings data, we excluded all individuals initially marked as an adult because uncertainty about their birth year precluded calculating key fitness-related traits (e.g. age at first breeding, longevity). We also excluded all individuals for which their sex at marking was unknown. Chicks can be sexed in the nest based on their morphometrics (Nicoll et al., 2003). We included all individuals marked as chicks irrespective of whether they were wild hatched or released because we have previously shown they have similar demography in early life (Nicoll et al., 2004). This resulted in 1,411 individual resighting histories available for analysis.

To explore inbreeding depression across the lifetimes of individual kestrels, we quantified the relationships between an individual’s level of inbreeding (*F*-coefficient) and various fitness-related demographic traits across its lifetime (Fig. 1). To do this, we used a structured statistical modelling approach implemented in R (R Core Team, 2024). We first fitted simple, univariate models to the data that included the various demographic traits as response variables and an individual’s *F*-coefficient as the predictor variable. Next, we fitted more complex models that included additional predictor variables known to influence the response variable of interest. We did this to ensure that any relationships between inbreeding and the response variable were unlikely to be confounded or obscured by additional ecological processes. We fitted separate models to the data for male and female kestrels, except for egg-to-fledgling survival models at the nest-scale because the sex of individual eggs was unknown. We checked all fitted models for convergence, overdispersion and collinearity between predictors. For the latter, we used the R package car (Fox and Weisberg, 2019).

Wherever necessary, we refitted models using the bobyqa optimizer, and standardised specific predictors by subtracting its mean (e.g. population density, individual age and its square).

<u>Parental kinship and egg survival:</u> We estimated the inbreeding coefficient of an egg by calculating the kinship coefficient of its parents (Kennedy et al., 2014). This generated a dataset of 624 nesting attempts for which the kinship of the parents was known, and pedigree depth was ≥2 generations. To quantify the relationship between parental kinship and egg-to-fledging survival, we fitted generalised linear mixed models (GLMMs) to the data using the lme4 package in R (Bates et al., 2015). We calculated the probability that an egg would survive to fledging in each nesting attempt using the cbind function (number of surviving eggs, number of eggs not producing a fledgling), and fitted GLMMs that included parental kinship as a continuous fixed effect, female ID as a random effect (we have repeat observations on some individuals across years), and assuming a binomial error distribution. More complex GLMMs included population density (standardised), nest type (box, cavity or tree), whether the nest was a first or second clutch, and female age (standardised linear and squared terms) as predictor variables (Burgess et al., 2011, Burgess et al., 2008, Cartwright et al., 2014a). We also included the *F*-coefficients of the parents as predictors in our more complex models to allow for inbreeding-related maternal or paternal effects (Richardson et al., 2004).

<u>Recruitment and longevity:</u> We quantified the relationship between inbreeding and three response variables based on the resightings data – the probability that a marked individual would be recruited into the breeding population (0 or 1); for recruited individuals, we calculated their age at first breeding (AFB); and we calculated individual longevity as the number of years between birth and an individual’s last resighting (see also Fig. 1). For the recruitment analyses, we fitted GLMs to the data in which recruitment probability was the response variable, an individual’s *F*-coefficient as a continuous predictor variable, and assuming binomial errors. For AFB, we fitted linear models with a similar structure assuming Gaussian errors. For the longevity data, we fitted Cox proportional hazard models (Cox, 1972). To do this, we censored individuals based on whether or not they were resighted in 2018/19. For more complex models of these response variables, we included cohort-level data on rainfall (wet/dry cyclone season OR raindays in the cyclone season) and population density (number of breeding pairs) for the birth year as these are known to affect cohort-survival in early life (Nicoll et al., 2003).

<u>Annual reproductive success (ARS):</u> We counted the total number of fledglings produced per breeding season by each adult bird recorded breeding (ARS). We then modelled ARS separately for each sex using GLMMs in which the total number of fledglings per breeding season per bird was the response variable, an individual’s *F*-coefficient was a continuous predictor variable, individual ID was a random effect, and assuming a negative binomial error distribution. We used this error distribution because initial analyses using Poisson errors showed significant overdispersion. For more complex models, we included population density (standardised) and individual age (standardised linear and square terms) as predictor variables because both are known to affect reproductive success (Burgess et al., 2008, Burgess et al., 2011, Cartwright et al., 2014b).

Differences in ARS in relation to inbreeding might occur for several reasons. First, some kestrel pairs form in a particular breeding season but do not lay any eggs. To see if this might be related to inbreeding, we fitted GLMMs to the data in which the probability of producing a clutch was the response variable, an individual’s *F*-coefficient was the predictor variable, individual ID was a random effect, and we assumed a binomial error distribution. Second, relatively inbred individuals might produce fewer eggs during a breeding season. To test this possibility, we calculated the total number of eggs produced per individual per breeding season, and fitted GLMMs to these data including an individual’s *F*-coefficient as the response variable, individual ID as a random effect and assuming a negative binomial error distribution. Third, inbred individuals might show reduced egg to fledgling survival rates. To test this possibility, we calculated the probability that an egg would survive to fledging in each breeding season using the cbind function (number of surviving eggs, number of eggs not producing a fledgling), and fitted GLMMs that included an individual’s *F*-coefficient as a continuous fixed effect, individual ID as a random effect, and assuming a binomial error distribution. For more complex models of these response variables, we included population density (standardised) and individual age (standardised linear and square terms). Separate models were fitted for males and females.

<u>Lifetime reproductive success (LRS):</u> We defined LRS as the total number of fledglings produced per individual across their entire lifetime. We then modelled LRS separately for each sex using GLMs in which an individual’s *F*-coefficient was a continuous predictor variable, and assuming a negative binomial error distribution. We included the number of breeding years in these models because LRS is known to be positively correlated with breeding lifespan in birds (Krüger and Lindström, 2003). Parameter estimates for the *F*-coefficient in these models provide direct estimates of lethal equivalents (Nietlisbach et al., 2019).

<u>Population viability:</u> We used an existing two-stage demographic model set-up in VORTEX to explore the implications of inbreeding depression for population growth and extinction risk (Nicoll et al., 2021). This model was developed to enable conservation scenarios to be explored, and requires details of reproductive and survival rates as input. To explore the impacts of inbreeding depression, we varied input parameters according to any fitness costs we found, and then ran the model for a range of inbreeding levels (details in the **Results**). For each inbreeding scenario, the model outputs the population growth rate (***r***) and the probability of extinction (***P_ext_***) over a 25- year period beginning with the population at carrying capacity (Nicoll et al., 2021).

## Results

### Inbreeding patterns

Inbreeding metrics are shown in Fig. 2. The distribution of inbreeding (*F*) coefficients is strongly left-skewed with most individuals being relatively outbred (*F* = 0) (Fig. 2A). Mean *F* = 0.068, and the range in *F* was 0-0.375. An *F* value of 0.25 indicates matings between full siblings or a parent and offspring, so some individuals showed very high levels of inbreeding. Mean levels of inbreeding increased over time before stabilising (Fig. 2B), but inbreeding varied between individuals across all study years (grey shaded area). Mean pedigree depth increased over time (Fig. 2C). These results show that the population has experienced significant levels of inbreeding.

**Figure 2.**
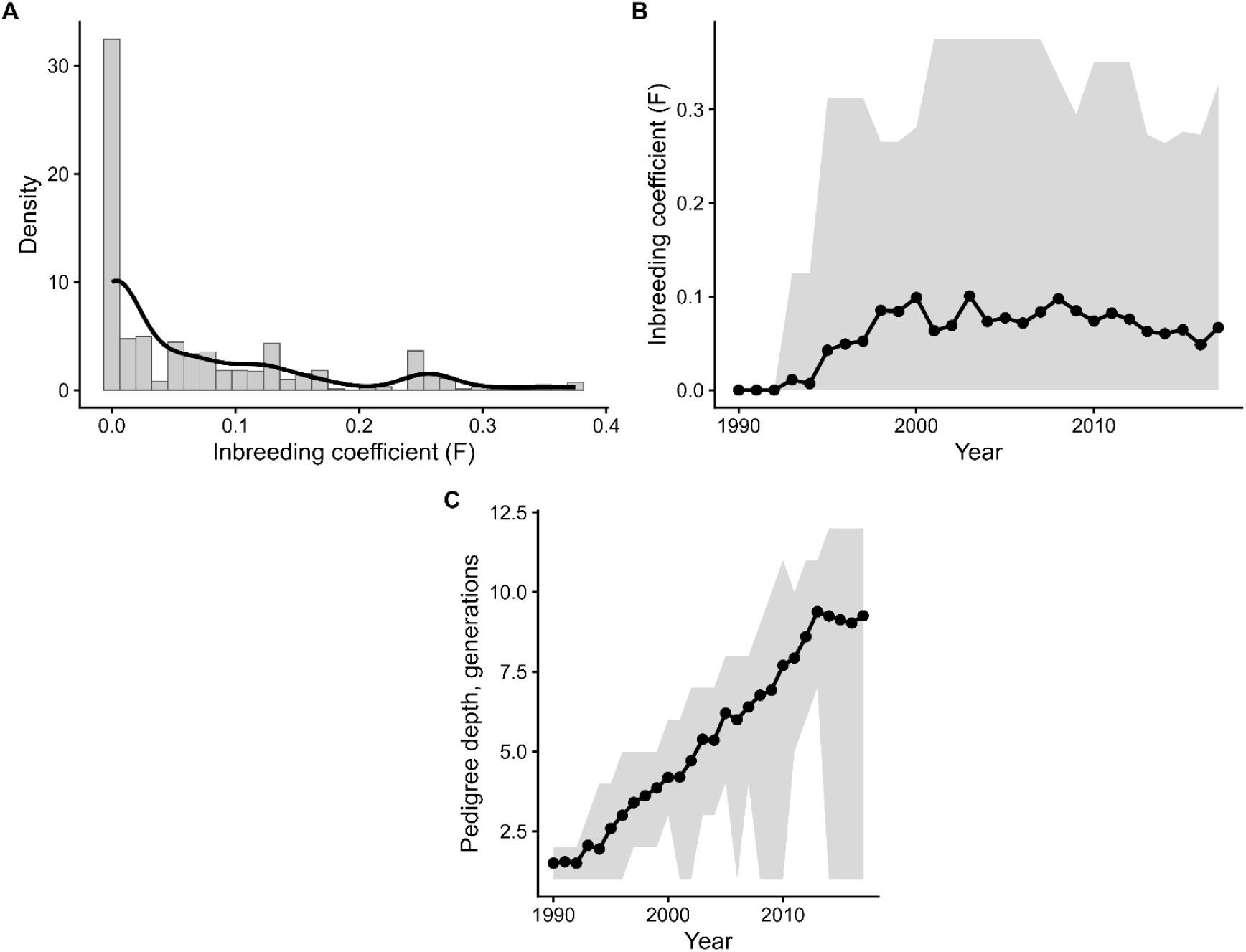
Patterns of inbreeding in the Mauritius kestrel population. Panel (**A**) shows a density histogram of inbreeding (*F*) coefficients. Panel (**B**) shows changes in inbreeding over time. The black points and solid black line show the mean annual *F*-value. The grey shaded area shows the range of individual *F*-values each year. Panel (**C**) shows changes in pedigree depth (generations) over time. The black points and solid black line show mean pedigree depth. The grey shaded area shows the range of individual pedigree depths each year.

### Parental kinship and egg survival

We found evidence of inbreeding depression in egg survival to fledging within individual nesting attempts. The relationship was statistically significant in the univariate model (β = −1.67±0.81, z-value = −2.05, p = 0.04), and in the more complex model (Table 1). The more complex model showed that egg survival to fledging decreased as population density increased, was significantly higher in first clutches, and showed a quadratic relationship with female age (increasing across early life age classes before declining into old age) (Table 1). We also found weak evidence of an inbreeding-related paternal effect (Table 1).

**Table 1.**
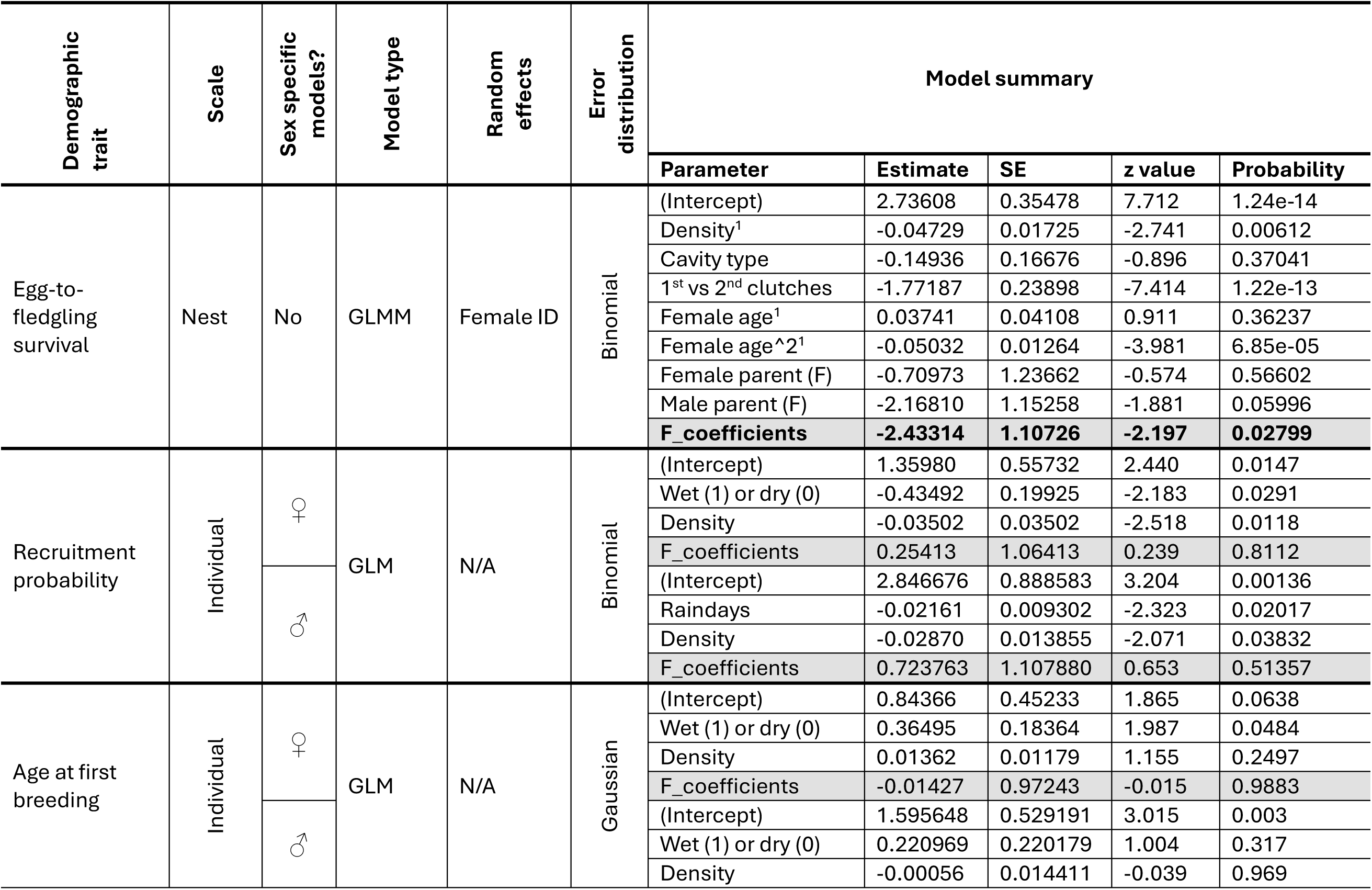
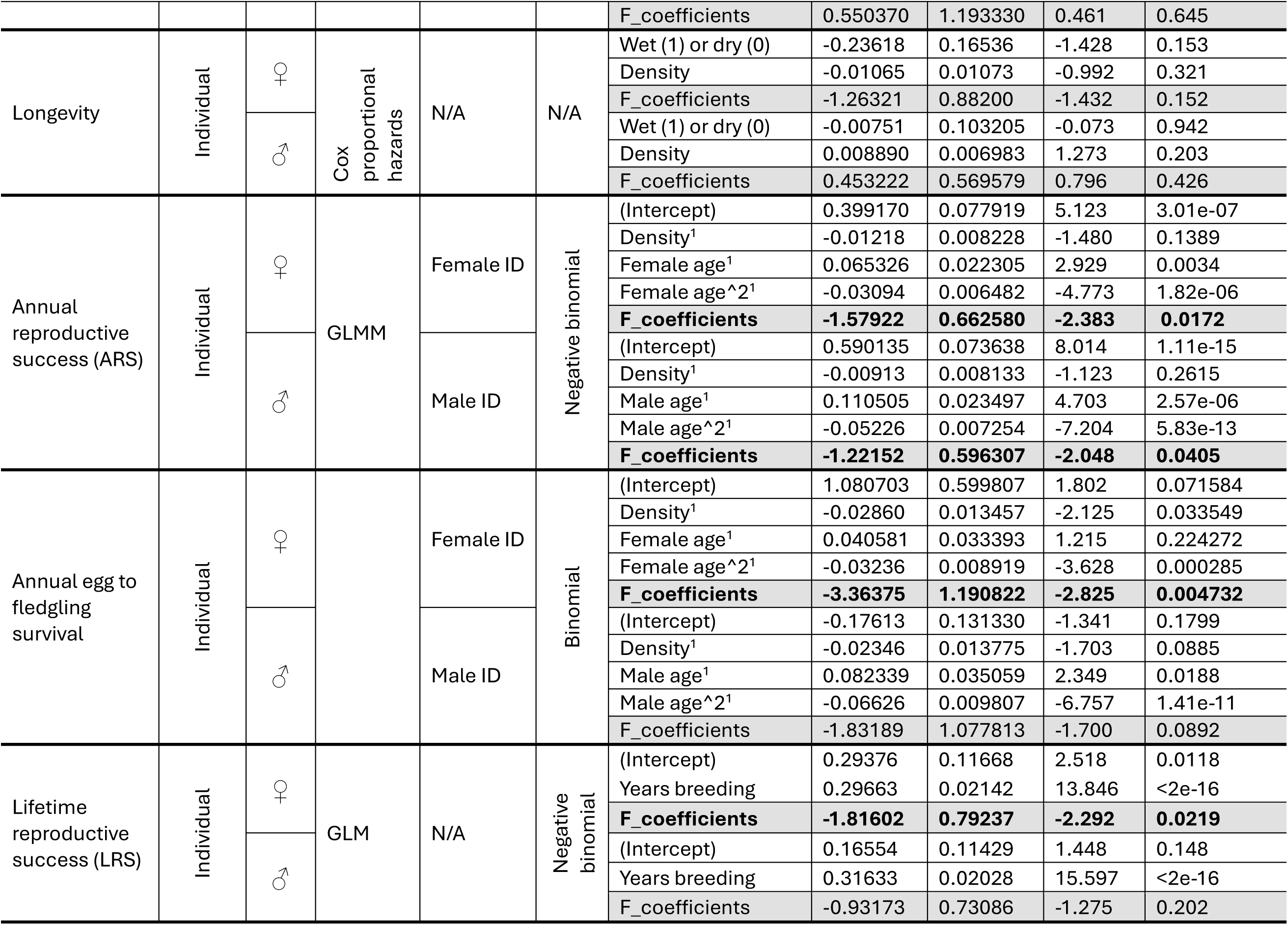
Statistical models quantifying the relationship between inbreeding (*F*) and fitness-related demographic traits. Inbreeding effects are shaded in grey; significant inbreeding effects are highlighted in bold. More detailed model descriptions are given in the text. ^1^Predictors were standardised.

### Recruitment and longevity

We found no evidence of significant inbreeding depression in the probability of recruitment, age at first breeding or longevity. The inbreeding (*F*) coefficient was not a significant predictor in the univariate or more complex models (Univariate models; Females; recruitment: β = 0.062±1.05, z-value = 0.06, p = 0.95; AFB: β = −0.28±0.99, z- value = −0.28, p = 0.78; longevity: β = −0.32±0.53, z-value = −0.61, p = 0.55; Males; recruitment: β = 0.50±1.09, z-value = 0.46, p = 0.65; AFB: β = 0.04±1.19, z-value = 0.03, p = 0.98; longevity: β = 0.53±0.56, z-value = 0.94, p = 0.35) (Table 1). The probability of recruitment was significantly lower for individuals fledging in years with frequent rainfall, and decreased as population density increased (Table 1). We also found that age at first breeding was significantly later in individuals fledging in years with frequent rainfall (Table 1). We found no significant environmental correlates of longevity (Table 1). For brevity, we only included models that distinguished wet/dry cyclone seasons, unless the model including raindays gave a contrasting result.

### Annual reproductive success (ARS)

ARS declined significantly with increasing levels of inbreeding in female and male kestrels (Fig. 3). The relationship was significant for both sexes in the univariate models (females: β = −1.56±0.65, z-value = −2.39, p = 0.017; males: β = −1.35±0.58, z-value = - 2.32, p = 0.02), and in the complex models (Table 1). For females, predicted ARS declines from 1.26 fledglings per year at *F* = 0 to 0.85 fledglings per year at *F* = 0.25, a decrease of ≈32%.

**Figure 3.**
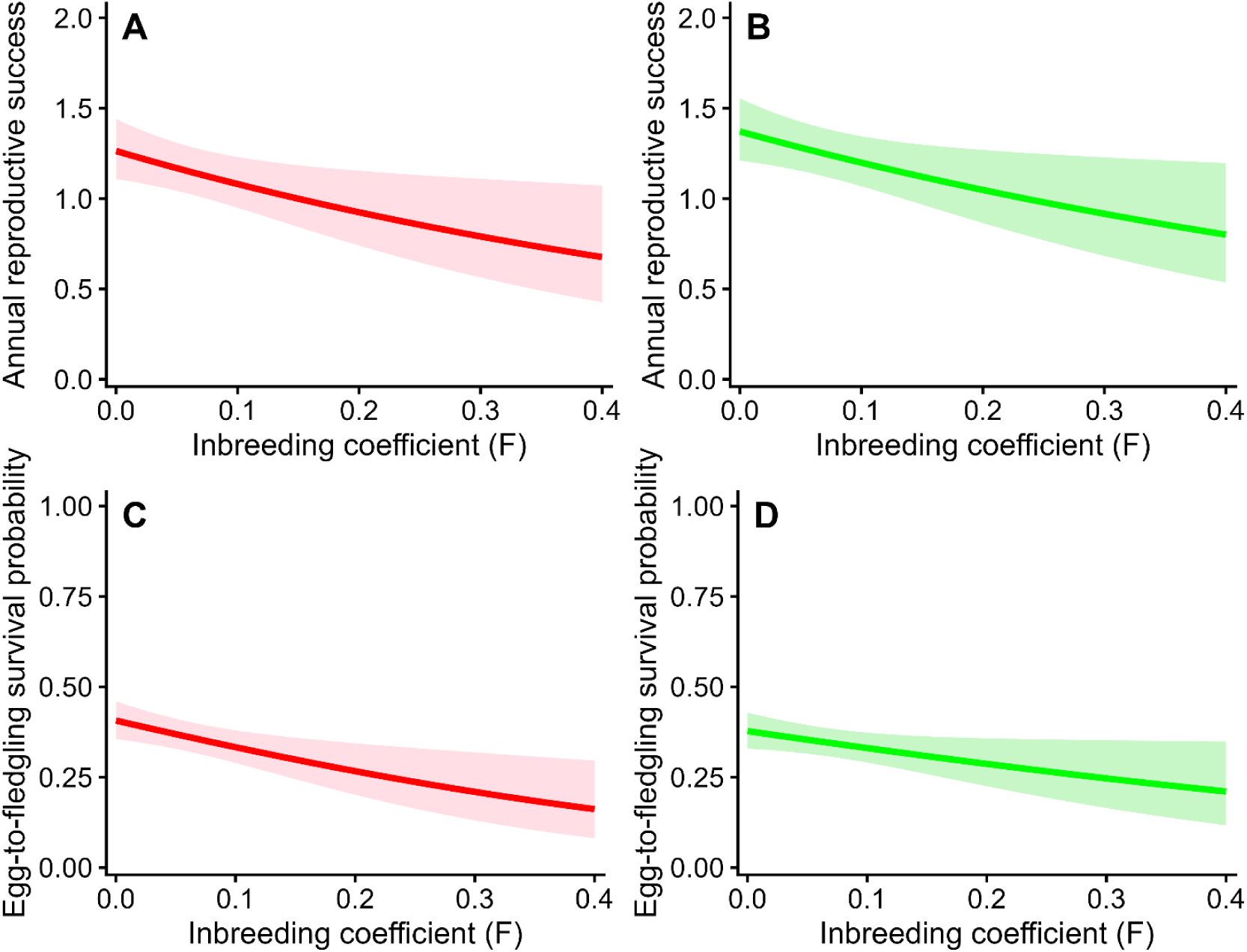
Inbreeding and annual reproductive success (ARS). Panels (**A**) and (**B**) show the relationships between ARS and inbreeding (*F*) coefficients for adult females (red) and adult males (green). Panels (**C**) and (**D**) show the relationships between annual egg-to-fledgling survival and inbreeding (*F*) coefficients for adult females (red) and adult males (green). Each plot shows the predicted values from univariate models (solid line) together with their 95% confidence intervals (paler ribbon). Model details are given in the text.

We found no clear evidence to suggest that this relationship might be because relatively inbred individuals were less likely to produce a clutch having formed a breeding pair (Univariate models; females: β = 1.61±5.22, z-value = 0.31, p = 0.76; males: β = - 0.17±4.7, z-value = −0.036, p = 0.97). Furthermore, there was no evidence that relatively inbred individuals produced fewer eggs per breeding season (Univariate models; females: β = 0.21±0.37, z-value = 0.59, p = 0.56; males: β = 0.38±0.33, z-value = 1.16, p = 0.25). We found similar patterns in the more complex models (Supplementary information; Table S1).

The annual probability of eggs surviving to fledging declined significantly with increasing inbreeding in female and male kestrels (Fig. 3). The relationship was significant for both sexes in the univariate models (females: β = −3.7±1.12, z-value = −2.82, p = 0.005; males: β = −2.06±1.02, z-value = −2.01, p = 0.04), and similar in the complex models (Table 1), although the trend was weaker in males.

### Lifetime reproductive success (LRS)

LRS declined significantly with increasing levels of inbreeding in females, while males showed a non-significant negative trend (Fig. 4; Table 1). These patterns were similar if individuals resighted in 2018/19 were excluded from analyses (females: β = −1.7±0.81, z-value = −2.09, p = 0.036; males: β = −0.93±0.73, z-value = −1.28, p = 0.2). Note that the parameter estimates associated with inbreeding (*F*-coefficients) in these models represent direct estimates of lethal equivalents (Nietlisbach et al., 2019). As expected, the number of breeding years was positively correlated with LRS in both sexes (Table 1).

**Figure 4.**
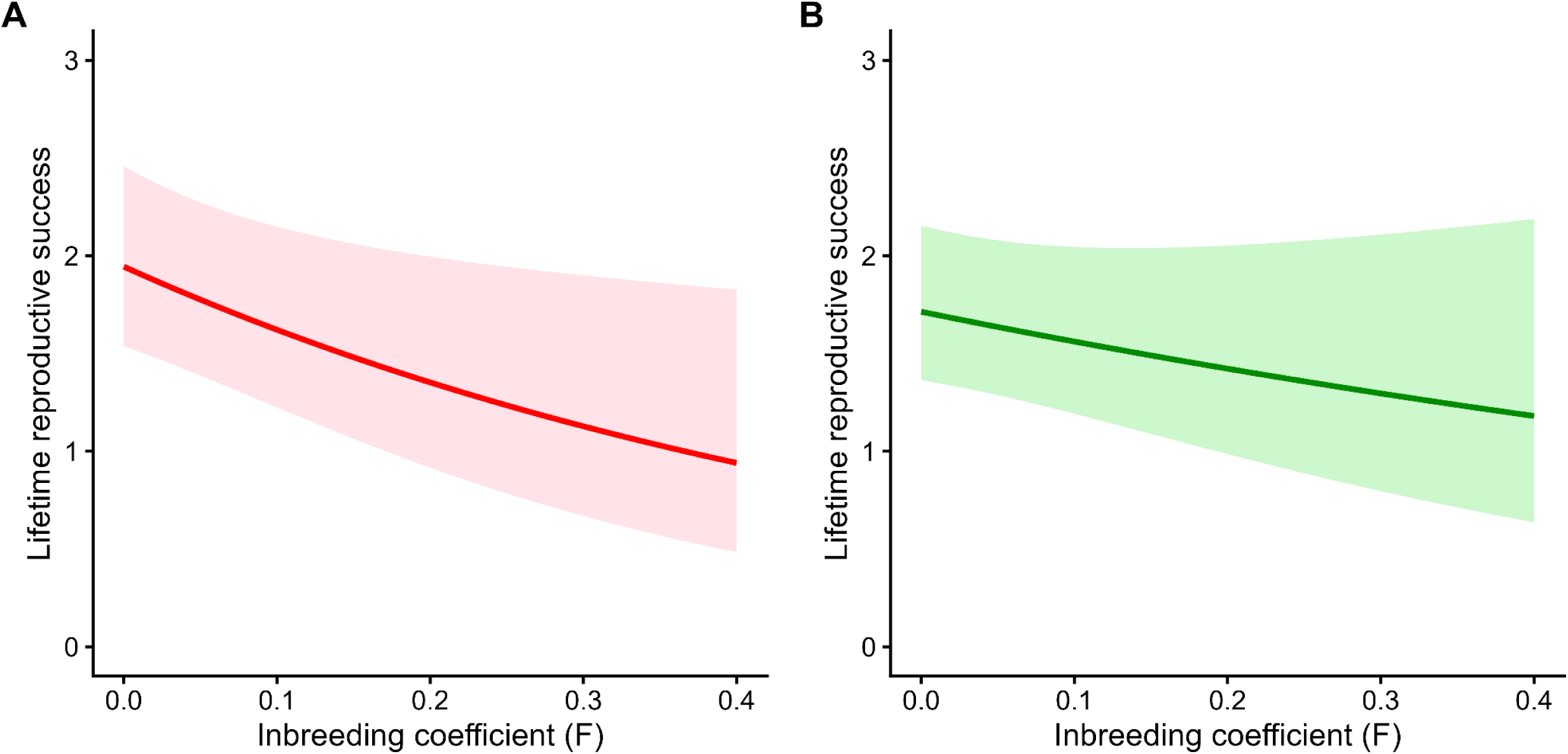
Inbreeding and lifetime reproductive success (LRS). Panels (**A**) and (**B**) show the relationships between LRS and inbreeding (*F*) coefficients for adult female (red) and adult males (green). Each plot shows the predicted values from univariate models (solid line) together with their 95% confidence intervals (paler ribbon). Model details are given in the text.

### Population viability

Our population model is female-only, and requires an estimate of fledgling production per female per year as input (reproductive rate) (Nicoll et al., 2021). This input is directly analogous to our measures of ARS (Fig. 3; Table 1). Therefore, we used the parameter estimates in our female ARS model to generate reproductive rates for mean population inbreeding ranging from *F* = 0 to *F* = 0.4, and ran our population model across this range for increments in *F* of 0.05. In this way, we explored the sensitivity of population growth and the probability of extinction to changes in mean levels of inbreeding within the kestrel population (central model scenarios). To incorporate uncertainty in our estimates of ARS in relation to inbreeding, we also estimated female ARS using the upper and lower 95% confidence limits of the parameter estimates, and ran our population model using these values across the same incremental range in *F* values (upper and lower 95% model scenarios).

The results are shown in Fig. 5. At observed levels of mean inbreeding (*F* = 0.068), simulated outcomes ranged from near stability under the upper 95% scenario to more negative growth and viability under the central and lower 95% scenarios (upper 95% scenario: *r* = −0.032, *P_ext_* = 0; central scenario: *r* = −0.085, *P_ext_* = 0.01; lower 95% scenario: *r* = −0.135, *P_ext_* = 0.1). Across the central model scenarios, increasing inbreeding was associated with progressively lower simulated population growth rates, and an increase in extinction risk. Results were sensitive to uncertainty in the fitted negative binomial model parameters. Under the upper 95% scenarios, population growth remained closer to stability across the range of *F* values considered and extinction risk remained negligible. In contrast, under the lower 95% scenarios, increasing *F* led to more strongly negative population growth and a marked increase in extinction risk.

**Figure 5.**
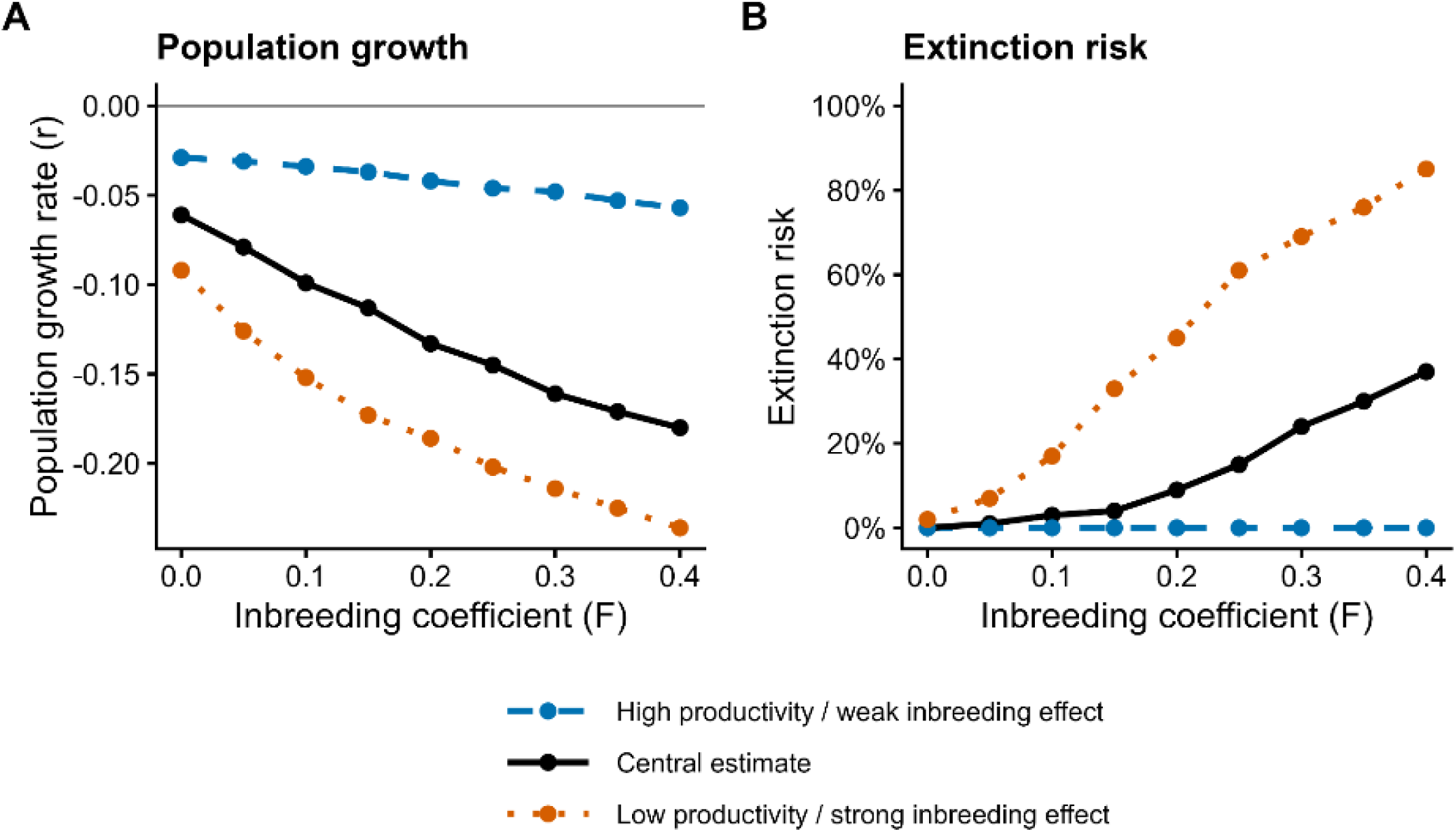
Simulated demographic consequences of reductions in annual reproductive success (ARS) in relation to mean inbreeding at the population-level. Lines show VORTEX simulation outcomes across a range of inbreeding coefficients (representing mean population-level inbreeding) using annual reproductive success values predicted from the negative binomial female ARS model (Table 1). The central scenario (black points) uses the fitted model estimates while upper (blue points) and lower (brown points) 95% model scenarios combine high baseline productivity with a weak inbreeding effect (upper 95% CIs of parameter estimates), and low baseline productivity with a strong inbreeding effect (lower CIs of parameter estimates), respectively. Panels show effects of inbreeding on (**A**) population growth rate (*r*) and (**B**) extinction risk (*P_ext_*).

## Discussion

Our analyses revealed significant fitness costs of inbreeding in Mauritius kestrels with important potential impacts on longer-term population viability. We demonstrated inbreeding depression in the egg-to-fledging survival rate within nests, and in the annual reproductive success of adult female and male birds driven by egg-to-fledgling survival rates – relatively inbred individuals had lower ARS and lower annual egg-to-fledgling survival rates compared with relatively outbred individuals. These differences translated into lower lifetime reproductive success in inbred females but not in males. Population modelling showed that inbreeding depression could reduce population growth and increase extinction risk, particularly if levels of inbreeding increased significantly in the future. In contrast, we found no evidence of inbreeding depression in demographic traits associated with survival (recruitment, age at first breeding, longevity).

The robustness of these results depends on the data and models on which they are based. Social pedigrees assume that social relationships define the relatedness between parents and offspring. This might not be the case if egg dumping or extra-pair fertilizations (EPFs) are common. We observed no cases of egg dumping over the study period (i.e. instances in which two females laid simultaneously in the same nest), and although we have no data on EPFs, these tend to be rare in falcons because within-pair copulations are frequent (Villarroel et al., 1998). Furthermore, we have whole genome data for a small sample of individuals included in our inbreeding depression study (Wang et al., 2026), and these data show significant correlations between genomic measures of inbreeding and *F* values derived from our social pedigree (Heterozygosity: *r* = −0.466, *p* = 0.029; Runs of homozygosity: *r* = 0.61, *p* = 0.0026; *df* = 20 in both cases). Taken together, this evidence suggests that the inbreeding coefficients calculated from our social pedigree provide a reasonable representation of variation in inbreeding between individual kestrels.

We found that inbreeding had increased over time in our study population before stabilising (Fig. 2b). This pattern most likely arises from the spatial arrangement of release sites and subsequent population expansion. Captive-reared birds were released at a series of sites (Burgess et al., 2008). Most (∼80%) of Mauritius kestrels disperse <2 km between their natal and breeding sites (Nevoux et al., 2013). As a result, three groups of release sites were separated by distances greater than this natal dispersal distance. As the population expanded, territories close to release sites were occupied first (Burgess et al., 2008), increasing the likelihood that individuals might breed with close relatives. This potentially explains why inbreeding increased during the early years. As the population expanded further, these release groups eventually overlapped, enabling pairs to form across release groups with individuals of lower relatedness. Over time, these release group lineages would become more thoroughly mixed. This likely explains why inbreeding eventually stabilised. This is an interesting example showing that the spatial arrangement of releases might affect the temporal development of inbreeding.

We used a structured statistical modelling framework to distinguish any potential inbreeding effects on demographic traits from other ecological processes that might amplify or mask them. To do so, we combined simple, univariate models with more complex models that included predictors known to be important drivers of variation in the demographic traits of interest. Our analyses drew on a substantial body of existing ecological knowledge on Mauritius kestrels (Nicoll et al., 2003, Nicoll et al., 2004, Burgess et al., 2011, Nevoux et al., 2011, Senapathi et al., 2011, Cartwright et al., 2014b, Cartwright et al., 2014a, Taylor et al., 2021). While our complex models revealed several important drivers of variation in demographic traits, there was generally good correspondence between the univariate and complex models in terms of inbreeding depression. While our analyses do not formally demonstrate causation, they do substantially reduce the likelihood that our inbreeding depression results can be explained by correlated ecological factors.

Our population modelling showed that population growth was negative and extinction risk slightly elevated at current levels of inbreeding (*F* = 0.068) when compared with an outbred population (*F* = 0) (Fig. 5). Our results suggest that these impacts will become more severe should inbreeding levels increase in the future, which is highly likely given ongoing genomic erosion (Wang et al., 2026). Differences in population growth and extinction risk between inbreeding scenarios should be regarded as relative rather than absolute because our model is sensitive to small changes in vital rates (Nicoll et al., 2021), and the inbreeding analysis was necessarily restricted to females with sufficient pedigree information. Nevertheless, the direction of the effect of inbreeding depression was consistent: increasing mean inbreeding at the population-level reduced predicted annual reproductive success and generally led to lower population growth and higher extinction risk. The results therefore suggest that inbreeding depression is unlikely to be merely a statistical signal in the reproductive success data, but represents an ecologically meaningful potential constraint on population viability. Similar results are emerging for other endangered birds (Armstrong et al., 2021, Trask et al., 2021). This suggests that inbreeding depression should be regarded as an important issue for the conservation management of Mauritius kestrels going forwards.

What mechanism might explain the inbreeding depression we observed? The most likely explanation is that inbreeding reduces egg-to-fledgling survival in the nest through a combination of reduced egg fertility, early embryo death in the egg, or lower chick survival. Hatching failure is common in threatened species (Marshall et al., 2023), and recent work has revealed high levels of early embryo death as a significant cause (Hemmings et al., 2012a, Morland et al., 2024). Studies have also shown that hatching failure and early death post-hatching can be related to inbreeding (Hemmings et al., 2012b, Hoeck et al., 2015, Kyriazis et al., 2026). Similar processes might explain the patterns we observed in the ARS of adult female and male kestrels, although a reduction in the quality of parental care provided by inbred birds might also play a role here (Richardson et al., 2004). Further work will be required to establish the timing and cause of death in the egg-to-fledgling period.

The fact that our analyses revealed significant inbreeding depression in Mauritius kestrels implies that deleterious, recessive mutations were not completely purged through the population bottleneck. This is confirmed by recent genomic analyses, which demonstrate significant and ongoing genomic erosion, and the incomplete purging of deleterious variants (Wang et al., 2026). Similar patterns are evident in several other endangered birds (Kennedy et al., 2014, Harrisson et al., 2019, Armstrong et al., 2021, Trask et al., 2021), suggesting incomplete purging might be common. While demographic recovery might offset this risk in some cases (Ahrens et al., 2026), the emerging evidence from wild birds suggests that genomic erosion and inbreeding are ongoing problems even in recovering populations (Cavill et al., 2024, Fontsere et al., 2025, Femerling et al., 2023). Our results suggest that Mauritius kestrels align with this avian pattern – although the population is recovering demographically, inbreeding and inbreeding depression remain a significant risk to population viability in the medium- to long-term.

Our analyses highlight the critical role played by long-term monitoring programmes in the context of endangered species recovery. The Mauritius kestrel programme will continue to provide key information on inbreeding, demographic change and population viability going forwards, and hence needs to be continued. Future options for managing inbreeding depression by increasing genetic diversity are limited because there are no alternative sources of birds for translocation (e.g. alternative populations or living collections). As a result, consideration needs to be given to interspecific hybridisation and genome engineering as possible future management options (Chan et al., 2019, Quilodran et al., 2020, Izquierdo-Aranega et al., 2025, van Oosterhout et al., 2025). This argues for a two-pronged approach to the conservation management of inbreeding and its consequences – ongoing population monitoring to assess inbreeding and its risks combined with a careful analysis of the risks and opportunities associated with future options to improve the genetic diversity of kestrels in the wild (Hogg et al., 2021).

In conclusion, we have demonstrated significant inbreeding depression in a recovering population of Mauritius kestrels with implications for population viability. This shows that inbreeding depression represents an ongoing risk to population recovery in the medium- to long-term, and therefore requires explicit consideration in conservation management planning. Our study contributes to a growing body of evidence showing similar patterns across a range of endangered bird species. This strongly suggests that demographic recovery is the start rather than the end of a journey from near extinction to a viable long-term population.

## Acknowledgements

The Mauritius Kestrel recovery programme has been supported by the Mauritian Wildlife Foundation, Standard Bank (Mauritius), Durrell Wildlife Conservation Trust, the Peregrine Fund, the Zoological Society of London, and the National Parks and Conservation Service (NPCS) (Government of Mauritius). The recovery programme and associated research would not have been possible without the cooperation of local landowners, and the dedication of numerous field biologists. Financial support to UK institutions was provided by Research England (including their E3 Fund), the Natural Environmental Research Council (NERC) and the Leverhulme Trust.

## Author contributions

Ken Norris conceived the ideas, designed the methodology, and led on the data analysis and interpretation, and the writing of the manuscript; Malcolm Nicoll and Carl Jones have led the research on Mauritius kestrels for many years, and contributed to data management, analysis and interpretation; Vikash Tatayah, Sion Henshaw and Nicolas Zuel lead the Mauritius kestrel conservation and monitoring programmes within Mauritius, and contributed to data management, analysis and interpretation; Kevin Ruhomaun is the Head of National Parks and Conservation Service, Government of Mauritius and has statutory oversight of and input to all the endangered bird conservation programmes, including Mauritius kestrels; Jim Groombridge, Hernán E. Morales, Cock van Oosterhout and Xuejing Wang lead on the Mauritius kestrel genomics work, and contributed to the data analysis and interpretation. All authors contributed critically to the manuscript and gave final approval for publication.

## Data availability statement

Data will be available through Zenodo http://dx.doi.org/10.5281/zenodo.21640455 (Norris 2026).

