## Supplementary figure and table for "Inbreeding depression and population viability in a recovering population of Mauritius kestrels"

### Supplementary Information

**Figure S1.** Temporal trends in the percentage of adult Mauritius kestrels associated with known territories or nests that were identified during the breeding season. Separate lines are shown for adult females, adult males and breeding pairs.

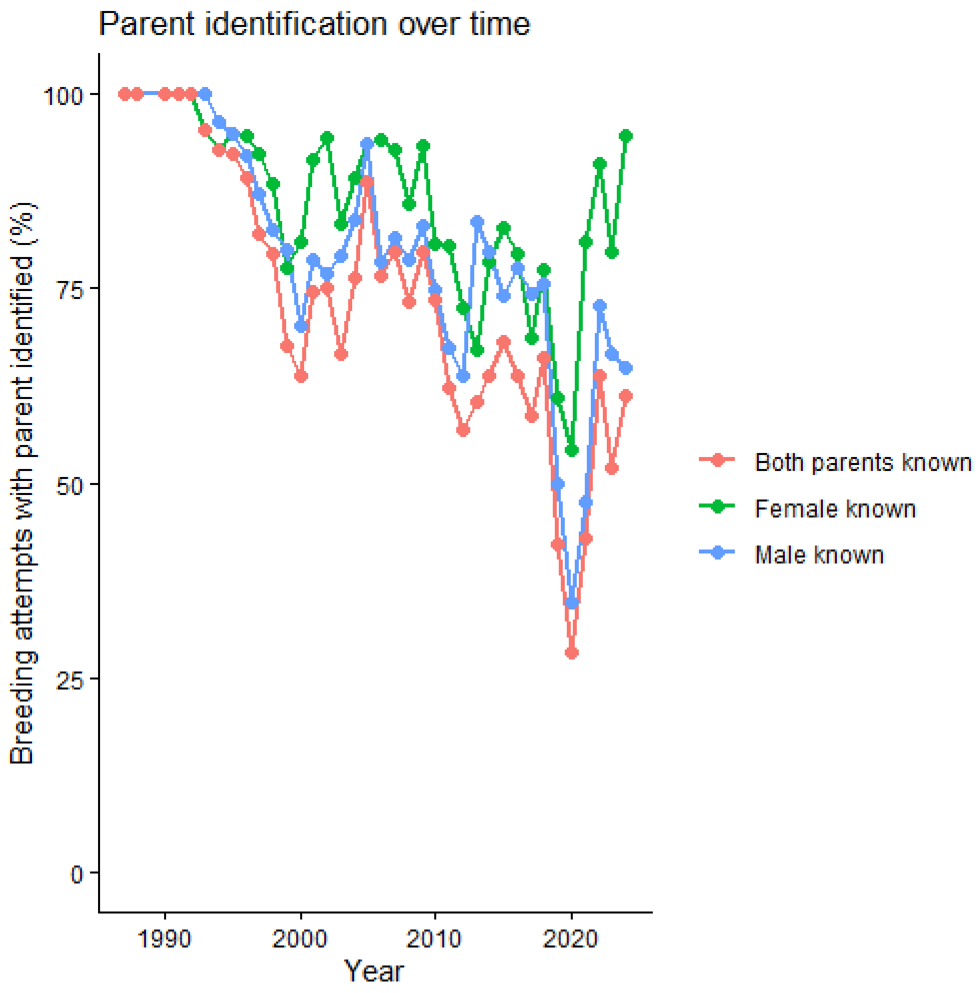

**Table S1.** Statistical models quantifying the relationship between inbreedng (*F*) and components of annual reproductive success. Inbreeding effects are shaded in grey; significant inbreeding effects are highlighted in bold. More detailed model desciptions are given in the text. [1]Standardised predictors.

| Demographic trait | Scale | Sex specific models? | Model type | Random effects | Error distribution | Model summary | | | | |
|---|---|---|---|---|---|---|---|---|---|---|
| | | | | | | **Parameter** | **Estimate** | **SE** | **z value** | **Probability** |
| Probability of producing eggs having formed a pair | Individual | ♀ | GLMM | Female ID | Binomial | (Intercept) | 4.85650 | 2.42529 | 2.002 | 0.04524 |
| | | | | | | Density[1] | -0.05747 | 0.04444 | -1.293 | 0.19592 |
| | | | | | | Female age[1] | 0.31604 | 0.09877 | 3.200 | 0.00138 |
| | | | | | | Female age^2[1] | -0.06402 | 0.02288 | -2.798 | 0.00514 |
| | | | | | | F_coefficients | 1.15078 | 3.75466 | 0.306 | 0.75923 |
| | | ♂ | | Male ID | | (Intercept) | 4.77840 | 0.88702 | 5.387 | 7.16e-08 |
| | | | | | | Density[1] | -0.08741 | 0.04392 | -1.990 | 0.0466 |
| | | | | | | Male age[1] | 0.45611 | 0.10674 | 4.273 | 1.93e-05 |
| | | | | | | Male age^2[1] | -0.10302 | 0.02441 | -4.220 | 2.44e-05 |
| | | | | | | F_coefficients | 0.13655 | 3.13866 | 0.044 | 0.9653 |
| Annual egg production | Individual | ♀ | GLMM | Female ID | Negative binomial | (Intercept) | 1.186683 | 0.048458 | 24.489 | < 2e-16 |
| | | | | | | Density[1] | 0.008525 | 0.005260 | 1.621 | 0.105082 |
| | | | | | | Female age[1] | 0.047856 | 0.013529 | 3.537 | 0.000404 |
| | | | | | | Female age^2[1] | -0.01293 | 0.003468 | -3.729 | 0.000193 |
| | | | | | | F_coefficients | 0.196922 | 0.388536 | 0.507 | 0.612273 |
| | | ♂ | | Male ID | | (Intercept) | 1.322912 | 0.046917 | 28.197 | < 2e-16 |
| | | | | | | Density[1] | 0.010038 | 0.005362 | 1.872 | 0.0612 |
| | | | | | | Male age[1] | 0.055549 | 0.055549 | 4.018 | 5.86e-05 |
| | | | | | | Male age^2[1] | -0.01556 | 0.003695 | -4.212 | 2.54e-05 |
| | | | | | | F_coefficients | 0.314166 | 0.342621 | 0.917 | 0.3592 |
